# Sources of Variability in Normative Cerebral [^18^F]FDG-PET imaging

**DOI:** 10.64898/2026.07.30.741623

**Authors:** Giordana Salvi de Souza, Guilherme Povala, Guilherme Garcia Schu Peixoto, Artur Martins Coutinho, Andrei Bieger, Mateus Rozalem-Aranha, Marco Antônio de Bastiani, Eduardo Rigon Zimmer, Wyllians Vendramini Borelli, Laura Willers de Souza

**Affiliations:** Graduate Program in Biological Sciences: Biochemistry, Universidade Federal do Rio Grande do Sul, Porto Alegre, Brazil; Department of Psychiatry, School of Medicine, University of Pittsburgh, Pittsburgh, PA, USA; Department of Research, Development and Innovation, masima: Macunaíma Soluções em Imagens Médicas, Porto Alegre, Brazil; Nuclear Medicine and PET/CT Service, Centro de Diagnósticos, Hospital Sírio-Libanês, São Paulo,Brazil; Laboratory of Nuclear Medicine (LIM 43), Institute of Radiology, Hospital das Clínicas, Faculdade de Medicina, Universidade de São Paulo (HC-FMUSP), São Paulo, Brazil; Neuroradiology Section, Department of Radiology, Hospital de la Santa Creu i Sant Pau, Biomedical Research Institute Sant Pau, Universitat Autònoma de Barcelona, Barcelona, Spain; Sant Pau Memory Unit, Hospital de la Santa Creu i Sant Pau, Institut de Recerca Sant Pau, Universitat Autònoma de Barcelona, Barcelona, Spain; Graduate Program in Biological Sciences: Pharmacology and Therapeutics, Universidade Federal do Rio Grande do Sul, Porto Alegre, Brazil; Hospital Moinhos de Vento, Porto Alegre, Brazil; Translational Neuroimaging Laboratory, The McGill University Research Centre for Studies in Aging, McGill University, 845 Sherbrooke Street West, H3A 2T5, Montreal, Quebec, Canada; Department of Pharmacology, Universidade Federal do Rio Grande do Sul, Porto Alegre, Brazil; Centro da Memória, Hospital Moinhos de Vento, Porto Alegre, Brazil

**Keywords:** quantitative analysis, PET imaging, cerebral metabolism, normative values

## Abstract

**Purpose:** Quantitative interpretation of brain [¹⁸F]FDG-PET increasingly relies on comparisons with normative datasets. However, normative values may be influenced by technical and biological factors, limiting their generalizability. We investigated the effects of scanner manufacturer, reference region, age, and sex on regional [¹⁸F]FDG uptake in cognitively normal (CN) adults and generated covariate-adjusted normative reference data.

**Methods:** A total of 449 CN participants from the Alzheimer’s Disease Neuroimaging Initiative (ADNI) were included. Regional SUVr were calculated using three reference regions (whole cerebellum, pons, cortical gray matter) and converted to Z-scores. Linear regression models were used to estimate standardized regression coefficients (β), and 10-fold cross-validation was performed to quantify the out-of-sample predictive contribution of each covariate using incremental explained variance (ΔR²).

**Results:** Scanner manufacturer introduced large, spatially structured biases. Compared with Siemens systems, GE and Philips scanners yielded lower Z-scores in frontal and medial temporal regions, with effect sizes approaching one standard deviation in selected regions (β up to −0.85). Age showed region-specific associations with subcortical nuclei, medial temporal structures, and the posterior cingulate cortex, and was the strongest biological predictor in cross-validation (ΔR²≈0.11). Sex effects were negligible (ΔR²<0.001). Cortical gray matter normalization minimized biological and technical confounding, and the AD meta-ROI demonstrated high robustness across manufacturers and normalization strategies.

**Conclusion:** Scanner manufacturer and age are the major sources of variance in brain [¹⁸F]FDG-PET quantification in CN subjects. Cortical gray matter provides the most stable reference region and supports harmonized, covariate-adjusted normative datasets for clinical and research applications.

## Introduction

[^18^F]fluorodeoxyglucose positron emission tomography ([^18^F]FDG-PET) is a well-established imaging technique used for evaluating cerebral glucose metabolism in research and clinical settings [1]. In these contexts, interpretation increasingly relies on quantitative comparisons, in which an individual’s standardized uptake value ratio (SUVr) is evaluated against normative distributions from cognitively normal (CN) subjects [2,3]. This normative approach, however, is highly sensitive to variability, particularly when reference data are insufficiently characterized. Such limitations are especially consequential for detecting subtle metabolic changes, where technical noise may obscure biologically meaningful effects [4].

Multiple methodological factors contribute to this heterogeneity in normative comparisons. Commercial software platforms (*e.g.*, CortexID, MIMNeuro, Scenium/syngo.via) rely on proprietary normative databases that differ in cohort composition, acquisition protocols, and validation strategies, limiting cross-platform generalization [5–7]. Even within large multicenter initiatives such as the Alzheimer’s Disease Neuroimaging Initiative (ADNI), a major source of reference data, standardization has primarily focused on acquisition protocols rather than on data harmonization across scanners and reconstruction methods. Consequently, residual inter-scanner differences persist. These effects, driven by scanner manufacturers, hardware generations, and reconstruction algorithms, may directly influence quantitative SUVr measurements and hinder the development of robust normative frameworks [4,8–11]. Additional methodological variability arises from the choice of reference region for normalization (*e.g.*, pons, whole cerebellum, cortical gray matter), as each region differs in susceptibility to partial volume effects and scanner-related artifacts [12,13].

Biological factors also influence brain glucose metabolism independently of disease. Sex-related differences in brain metabolism have likewise been reported and are thought to reflect hormonal influences and sex-specific network organization [14–16]. Also, aging, for instance, produces widespread but regionally specific declines in brain [^18^F]FDG uptake, particularly in cortical areas such as temporoparietal and frontal regions [15,16]. Accounting for these covariates is essential for developing normative models that accurately capture interindividual variability in healthy populations. However, although normative [¹⁸F]FDG-PET datasets are available, the combined influence of scanner manufacturer, scanner technology, reference region selection, and biological factors remains underexplored. Current evidence has predominantly examined these factors in isolation. For instance, reconstruction protocols and scanner differences are known to introduce significant variability in quantitative measures [17,18], while the choice of reference region, such as the pons versus the whole cerebellum, can substantially impact results and interact with biological variables [19].

With this in mind, we leveraged [^18^F]FDG-PET data from CN older adults from the multicenter ADNI cohort to quantify and compare the effects of age, sex, scanner manufacturer, and scanner technology on regional [^18^F]FDG-PET measures, assess whether reference region selection modulates these effects, and generate adjusted normative scores incorporating these predictors.

## Material and methods

### Subjects

Data were obtained from the ADNI cohort (http://adni.loni.usc.edu), an international, multicenter longitudinal study involving 50 sites across the United States and Canada. Participants classified as cognitively normal (CN) were selected across the ADNI1, ADNI GO, ADNI2, and ADNI3 phases. CN status was defined by Mini-Mental State Examination (MMSE) scores of 24–30, a global Clinical Dementia Rating (CDR) of 0, and no clinical diagnosis for mild cognitive impairment or Alzheimer’s disease. Demographic, clinical, and cognitive data were collected from all participants. Eligibility criteria are detailed in the ADNI study protocol (www.adni-info.org/Scientists/ADNIStudyProcedures). Written informed consent was obtained at all sites in accordance with local ethics regulations.

### ADNI PET scanning procedures

Baseline brain [^18^F]FDG-PET scans were acquired following standardized ADNI protocols. Participants fasted for at least 4 hours before receiving an intravenous injection of approximately 185 MBq of [^18^F]FDG. Image acquisition started 30 minutes post-injection and lasted approximately 30 minutes (30-60 time window).

Due to the multicenter and multi-phase nature of ADNI, PET data used in this study were obtained using a wide range of scanner manufacturers (GE Healthcare, Siemens Healthineers, Philips Healthcare) and different models, including GE Advance, GE Discovery LS, GE Discovery RX, GE Discovery ST, GE Discovery 600, GE Discovery STE/VCT, Siemens EXACT, Siemens HR+, Siemens HRRT, Siemens ACCEL, Siemens ACCEL/EXACT, Siemens Biograph, Siemens Biograph (1023/1024), Siemens Biograph HiRes (1080), Siemens BioGraph HiRez, Siemens Biograph TruePoint, Siemens Biograph mCT, Philips Allegro, Philips Allegro-Neuro, Philips Gemini, Philips Gemini GLX, and Philips Gemini TF. Despite these model differences, all scans adhered to the same ADNI guidelines for participant preparation, injection, and acquisition time, ensuring comparable baseline conditions across sites. Assuming unbiased image reconstruction, summing dynamic frames produces results equivalent to a static acquisition of the same duration.

### Image processing

Raw T1-weighted MRI data (3T) scans were acquired as part of the ADNI acquisition protocols. Detailed information regarding MRI acquisition and preprocessing is available at http://adni.loni.usc.edu/methods/mri-tool/mri-analysis. Linear and non-linear image registration was performed using Advanced Normalization Tools (ANTs) [20] to align images to the ADNI template space [21]. All registrations underwent visual quality control to ensure accurate alignment.

Preprocessed [^18^F]FDG-PET data from the ADNI database were downloaded. Further details about the ADNI PET data acquisition protocol and data pre-processing are available at http://adni.loni.usc.edu/methods/pet-analysis-method/pet-analysis/. PET images were first rigidly registered to each subject’s T1-weighted MRI using ANTs. Subsequently, linear and non-linear T1-derived transformations were applied to warp PET images into ADNI stereotaxic space, ensuring anatomical correspondence across participants. PET images were then spatially smoothed with an 8 mm full-width at half maximum (FWHM) Gaussian kernel. All normalized images underwent visual quality control.

[^18^F]FDG SUV was measured in brain regions defined by the Desikan-Killiany-Tourville (DKT) atlas [22], including cortical lobes and basal ganglia. In addition to individual regions, we constructed a composite meta-region of interest (meta-ROI) averaging SUVr across established AD-vulnerable areas: bilateral angular gyri, temporal cortex (middle and inferior temporal gyri), and posterior cingulate cortex, as previously described by Landau *et al.* [23]. SUVr were calculated using three reference regions: cortical gray matter, pons, and whole cerebellum.

### Normative statistics

For each brain region, lobe, and reference region normalization strategy, SUVr values were transformed into Z-scores to facilitate comparison across regions and individuals. Z-scores were calculated using the SUVr values distribution. To provide clinically applicable reference data, we subsequently generated stratified normative tables (normograms) for all brain regions across the age range of 20 to 90 years, accounting for the key predictors identified in our models.

### Sensitivity Analysis

To evaluate whether the inclusion of participants with underlying amyloid pathology influenced associations, all primary analyses were repeated in the subset of participants classified as amyloid-negative at baseline. Amyloid status was determined using available AV45 PET or CSF Aβ42 data (UPENN Luminex assay), applying established ADNI thresholds: amyloid-positive was defined as whole-cerebellum-normalized [^18^F]AV45 SUVR ≥ 1.11 [24] or CSF Aβ42 < 192 pg/mL. [^18^F]AV45 PET was used as the primary biomarker; CSF Aβ42 served as fallback when PET data were unavailable. Participants meeting neither positivity criterion were classified as amyloid-negative.

### Statistical Analysis

Linear regression models were used to predict Z-score values for each brain region. Models were fitted separately for all 83 individual DKT regions, for each cortical lobe (frontal, parietal, temporal, and occipital), and for the AD meta-ROI. Age, sex, and scanner manufacturer (GE, Philips, Siemens) were included as covariates, with Siemens as the reference manufacturer because of its larger sample size. P-values were adjusted for multiple comparisons using the Benjamini-Hochberg false discovery rate (FDR) method with a threshold of q < 0.05. The Akaike Information Criterion (AIC), Bayesian Information Criterion (BIC), and coefficient of determination (R²) were evaluated for each model.

To evaluate the influence of scanner technology beyond the manufacturer, scanners were also classified into functional groups based on manufacturer, modality, and time-of-flight (TOF) capability (Supplementary Table 1). These groups included GE/Siemens/Philips standalone PET, GE/Siemens/Philips PET/CT without TOF, GE/Siemens/Philips PET probable with TOF, and GE/Siemens/Philips modern PET with TOF. The scanner group was included as a covariate in additional models to account for variability introduced by acquisition protocols, reconstruction algorithms, and other technical factors.

To assess model robustness and reduce overfitting, 10-fold cross-validation was performed as described by Potvin *et al.* [25]. Out-of-sample predictive contribution (ΔR²), defined as the difference in R² between the full model and the model without the predictor of interest, was calculated for each predictor by comparing full and reduced models in held-out data. All statistical analyses were performed in R (*version* 4.3; R Foundation for Statistical Computing, Vienna, Austria).

## Results

### Demographics

A total of 449 CN participants were included and divided into three age groups (≤65, 65–75, and ≥75 years). The groups did not differ in sex distribution, education, MMSE scores, or APOE4 frequency. Race varied across age categories, with a higher proportion of white participants in groups over 65 years old (*p = 0.002*). Imaging was performed predominantly on Siemens PET systems (55%), followed by GE (29%) and Philips (16%). Detailed demographic characteristics are shown in Table 1.

**Table 1.** Sample demographics and clinical characteristics.

|  | 65 years or below<br>(n = 21) | 65 to 75 years old<br>(n = 247) | 75 years or above<br>(n = 181) | *p-value* |
| --- | --- | --- | --- | --- |
| Age | 63 (2)† | 70 (3)† | 80 (3)† | < 0.001 |
| Female sex | 10 (48%) | 126 (51%) | 90 (50%) | 0.9 |
| Education (years) | 17 (2) | 16 (3) | 16 (3) | 0.5 |
| MMSE | 29 (1) | 29 (1) | 29 (1) | 0.8 |
| CDR | 0.1 (0.2) | 0.04 (0.14) | 0.09 (0.27) | 0.08 |
| APOE4 carrier | 0 | 6 (2.4%) | 5 (2.8%) | 0.9 |
| ICV (mm<sup>3</sup>) | 1,518 (188) | 1,503 (153) | 1,522 (162) | 0.5 |
| Amyloid<br>(neg/pos/miss), % | 87.0/4.3/8.7 | 65.4/18.7/15.9 | 55.0/25.6/19.4 | 0.017 |
| *Race* |  |  |  |  |
| White | 14 (67%)‡ | 227 (92%) | 166 (92%) | 0.002 |
| Black | 1 (4.8%) | 15 (6.1%) | 10 (5.5%) |  |
| Other races | 6 (29%)* | 5 (2.0%) | 5 (2.8%) |  |
| *Scanner* |  |  |  |  |
| GE | 5 (24%) | 82 (33%) | 52 (29%) | 0.6 |
| Philips | 2 (9.5%) | 31 (13%) | 29 (16%) |  |
| Siemens | 14 (67%) | 134 (54%) | 100 (55%) |  |
| *Image acquisition date* |  |  |  |  |
| 2005 - 2009 | 5 (24%) | 51 (21%) | 51 (28%) | 0.2 |
| 2010 - 2015 | 16 (76%) | 196 (79%) | 130 (72%) |  |

**Footnote:** †All pairwise comparisons for age were significant ( $p < 0.001$ ). ‡ The $\leq 65$ years group also differed significantly from both older groups for White and Other races ( $p < 0.001$ ). Amyloid positivity differed significantly across age groups after excluding participants with missing amyloid status (Pearson $\chi^2(2) = 8.18$ , $p = 0.017$ ; Cramér's $V = 0.15$ ). **Abbreviations:** MMSE, Mini-Mental State Examination; CDR, Clinical Dementia Rating; APOE4, apolipoprotein E $\epsilon 4$ allele; ICV, intracranial volume.

### Major effects of Lobar and Meta-ROI Z-Scores

To quantify the effects of biological and technical variables, we first assessed their influence at the lobar level and for an AD meta-ROI. Figure 1 shows the pattern of significant standardized regression coefficients (β) for age, sex, and scanner manufacturer across the three reference regions. β represents the variation in Z-score (SD units) associated with each predictor for scanner manufacturer; coefficients reflect differences relative to Siemens.

**Fig. 1.**
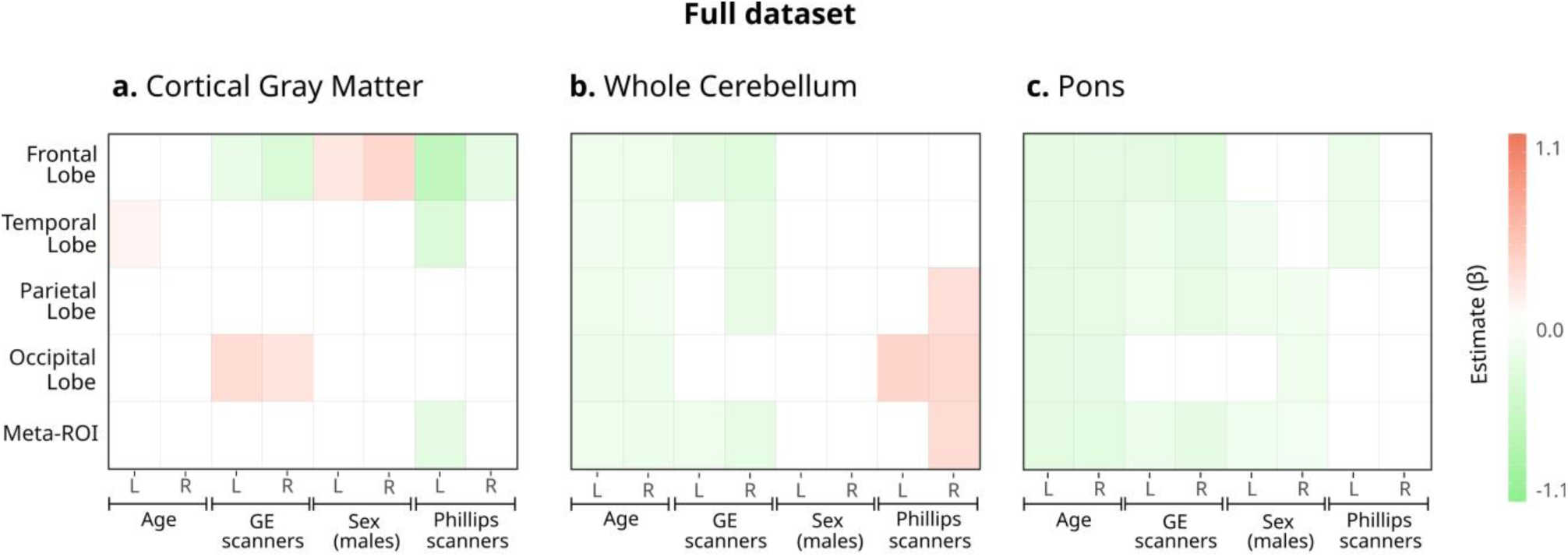
Heatmap of standardized regression coefficients (β) for age, sex, and scanner-related predictors on lobar and AD meta-ROI [¹⁸F]FDG Z-scores, stratified by hemisphere (left/right). A. Represents normalization by cortical gray matter; B. Represents normalization by whole cerebellum; C. Represents normalization by pons. Only FDR-corrected significant coefficients (p < 0.05) are displayed; nonsignificant effects are omitted.

Normalization using the whole cerebellum (Fig. 1B) or the pons (Fig. 1C) introduced pronounced, widespread negative age associations across all lobes (*p<0.001*). In contrast, under cortical gray matter normalization (Fig. 1A), age and sex effects were minimal. Scanner manufacturer (GE and Philips *vs*. Siemens) showed significant but regionally specific effects.

The lobar analysis revealed distinct patterns. The frontal lobes showed the strongest and most consistent associations with non-biological factors, being significantly influenced by sex and scanner manufacturer. The occipital lobes were selectively associated with the scanner manufacturer only. In contrast, the parietal and temporal lobes exhibited minimal sensitivity to these confounders. The AD meta-ROI demonstrated exceptional robustness, with negligible variance explained by any predictor across all reference regions (adjusted R² ≤ 0.03), indicating that composite measures are less susceptible to this technical and biological variability. Detailed coefficients for the cortical gray matter normalization are presented in Table 2. Results for pons and whole cerebellum normalization are included in Supplementary Tables 2–3.

**Table 2.** Standardized regression coefficients (β) for lobe-level and meta-ROI [¹⁸F]FDG Z-score normalized to cortical gray matter.

|  | Intercept | Age | Sex<br>(male) | Scanner<br>manufacturer<br>(reference<br>Siemens) = |  | Total<br>R<sup>2</sup> | Total<br>adjusted<br>R<sup>2</sup> | AIC | BIC |
| --- | --- | --- | --- | --- | --- | --- | --- | --- | --- |
|  |  |  |  | GE | Philips |  |  |  |  |
| Right |  |  |  |  |  |  |  |  |  |
| Meta-ROI | -0.04 | 0.04 | 0.13 | -0.08 | 0.02 | 0.01 | 0 | 1263.57 | 1288.22 |
| Frontal lobe | -0.01 | 0.02 | **0.45\*\*\*** | **-0.53\*\*\*** | **-0.39\*** | 0.12 | 0.11 | 1228.34 | 1252.98 |
| Temporal lobe | -0.06 | 0.08 | 0.19 | -0.03 | -0.21 | 0.02 | 0.01 | 1260.70 | 1285.34 |
| Parietal lobe | 0.09 | 0.1 | -0.06 | -0.17 | -0.01 | 0.02 | 0.01 | 1276.91 | 1301.55 |
| Occipital lobe | -0.05 | -0.03 | -0.12 | **0.3\*** | 0.15 | 0.02 | 0.01 | 1274.51 | 1299.15 |
| Left |  |  |  |  |  |  |  |  |  |
| Meta-ROI | -0.01 | 0.08 | -0.01 | 0.21 | **-0.38\*** | 0.04 | 0.03 | 1268.59 | 1293.23 |
| Frontal lobe | 0.1 | -0.01 | **0.26\*** | **-0.33\*** | **-0.88\*\*\*** | 0.11 | 0.11 | 1231.28 | 1255.92 |
| Temporal lobe | -0.02 | **0.12\*** | 0.07 | 0.19 | **-0.51\*** | 0.06 | 0.05 | 1257.35 | 1281.99 |
| Parietal lobe | 0.05 | 0.01 | -0.17 | 0.21 | -0.21 | 0.02 | 0.02 | 1273.90 | 1298.54 |
| Occipital lobe | -0.17 | -0.07 | 0.03 | **0.39\*** | 0.25 | 0.04 | 0.03 | 1268.19 | 1292.83 |

\*\*\*\* $p < 0.001$ ; \*\* $p < 0.01$ ; \* $p < 0.05$ . **Abbreviations:** AIC, Akaike Information Criterion; BIC, Bayesian Information Criterion; Meta-ROI, meta-region of interest comprising AD-vulnerable areas.

### Predictors of Individual Brain Regions

To further quantify and compare the effects of these variables at a finer anatomical scale, we performed region-wise analyses. Figure 2 presents a heatmap of significant effects across all brain regions for the three reference schemes, revealing distinct spatial patterns.

**Fig. 2.**
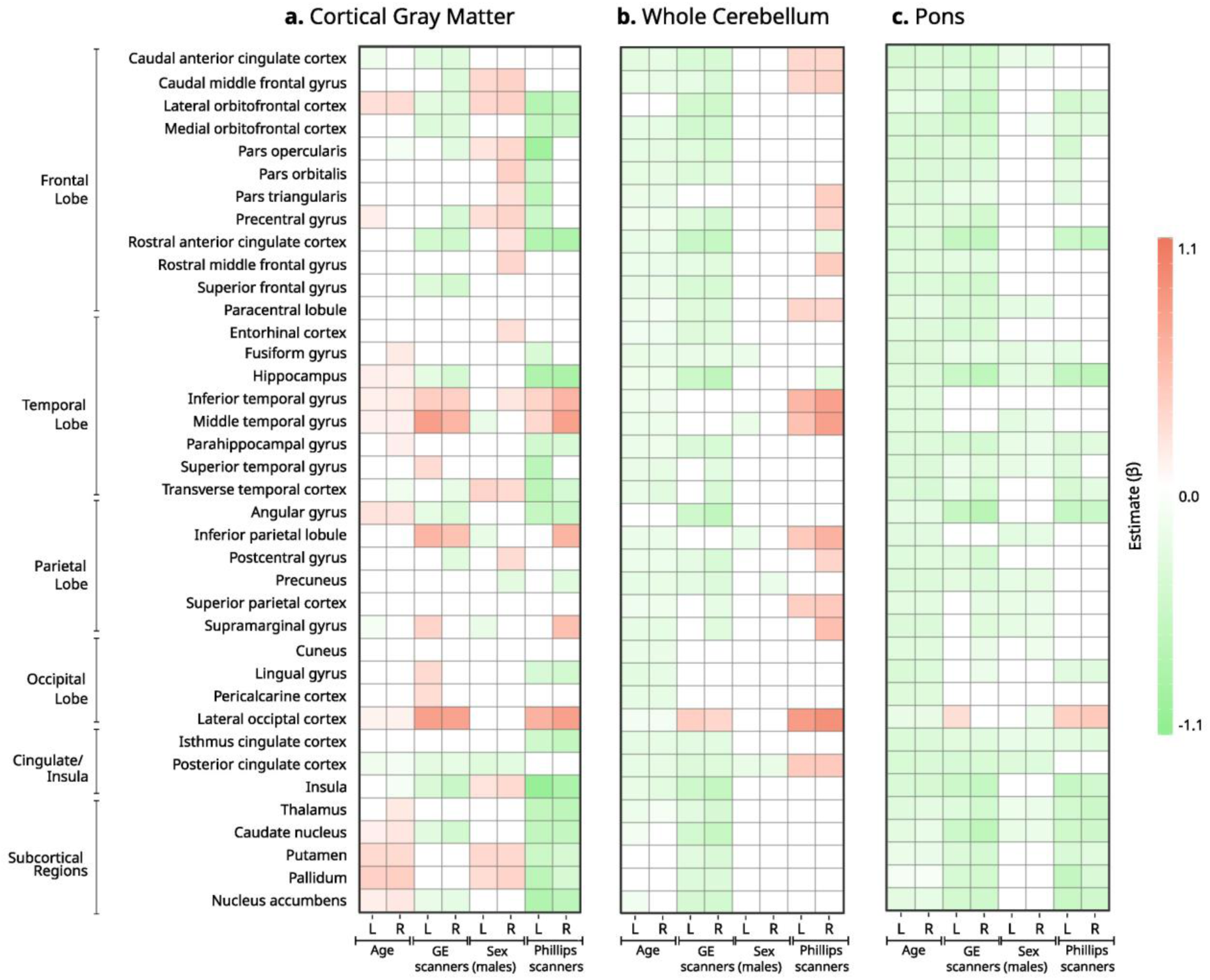
Heatmap of standardized regression coefficients (β) for age, sex, and scanner-related predictors on regional [¹⁸F]FDG Z-scores across brain regions. Stratified by hemisphere (left/right). A. Represents normalization by cortical gray matter; B. Represents normalization by whole cerebellum; C. Represents normalization by pons. Only FDR-corrected significant coefficients (*p < 0.05*) are displayed; nonsignificant effects are omitted.

Cortical gray matter normalization minimized scanner-related variability and reduced the spatial extent of age- and sex-related effects. For conciseness, Table 3 presents standardized coefficients (β) for a curated set of brain regions that best exemplify the major spatial patterns identified in the full analysis (see complete results for all 83 regions in Supplementary Table 4). The selected regions represent: (1) key nodes of the default mode network; (2) a medial temporal lobe structure; (3) frontal association cortices where scanner effects were pronounced; (4) subcortical nuclei with strong age and sex associations; and (5) a primary sensory region for contrast. Descriptive results for the results for pons and whole cerebellum normalization are included in Supplementary Tables 2–3. Figure 3 illustrates the cortical distribution of age, sex, and scanner effects using cortical gray matter normalization.

**Table 3.** Standardized regression coefficients (β) for regional [¹⁸F]FDG Z-score using cortical gray matter as the reference region.

|  | Intercept | Age | Sex (male) | Scanner manufacturer<br>(reference = Siemens) |  | Total R<sup>2</sup> | Total R<sup>2</sup> adjusted | AIC | BIC |
| --- | --- | --- | --- | --- | --- | --- | --- | --- | --- |
|  |  |  |  | GE | Philips |  |  |  |  |
| *Right* |  |  |  |  |  |  |  |  |  |
| Angular gyrus | 0.12 | **0.23\*\*\*** | 0.16 | **-0.38\*\*** | **-0.55\*\*** | 0.11 | 0.1 | 1232.08 | 1256.72 |
| Caudate | **0.23\*** | **0.2\*\*\*** | -0.01 | -<br>**0.46\*\*\*** | **-0.62\*\*\*** | 0.1 | 0.1 | 1235.45 | 1260.09 |
| Hippocampus | **0.28\*\*** | **0.12\*** | -0.06 | -<br>**0.43\*\*\*** | **-0.85\*\*\*** | 0.11 | 0.1 | 1234.70 | 1259.34 |
| Medial orbitofrontal | 0.15 | -0.03 | 0.06 | **-0.35\*** | **-0.54\*\*** | 0.05 | 0.04 | 1263.57 | 1288.22 |
| Pallidum | -0.06 | **0.39\*\*\*** | **0.34\*\*** | -0.16 | **-0.43\*** | 0.22 | 0.21 | 1176.37 | 1201.02 |
| Posterior cingulate | **0.2\*** | **-0.12\*** | **-0.3\*** | **-0.26\*** | 0.25 | 0.07 | 0.06 | 1253.81 | 1278.45 |
| Precuneus | **0.24\*** | -0.05 | **-0.28\*** | -0.19 | **-0.33\*** | 0.04 | 0.03 | 1267.91 | 1292.55 |
| Putamen | -0.04 | **0.3\*\*\*** | **0.31\*** | -0.18 | **-0.43\*** | 0.14 | 0.14 | 1215.13 | 1239.77 |
| Superior frontal | 0.08 | -0.01 | 0.16 | -<br>**0.46\*\*\*** | -0.09 | 0.05 | 0.04 | 1263.13 | 1287.77 |
| Thalamus | 0.13 | **0.18\*\*** | 0.03 | -0.17 | **-0.68\*\*\*** | 0.08 | 0.07 | 1247.81 | 1272.46 |
| *Left* |  |  |  |  |  |  |  |  |  |
| Angular gyrus | 0.1 | **0.22\*\*\*** | 0.13 | **-0.25\*** | **-0.6\*\*\*** | 0.1 | 0.09 | 1240.14 | 1264.78 |
| Caudate | **0.21\*** | **0.12\*** | -0.06 | **-0.31\*** | **-0.63\*\*\*** | 0.06 | 0.05 | 1256.18 | 1280.82 |
| Hippocampus | **0.24\*** | **0.13\*** | -0.08 | **-0.28\*** | **-0.8\*\*\*** | 0.09 | 0.08 | 1244.79 | 1269.43 |
| Medial orbitofrontal | 0.14 | -0.02 | 0.11 | **-0.37\*** | **-0.62\*\*\*** | 0.06 | 0.05 | 1257.24 | 1281.89 |
| Pallidum | -0.01 | **0.37\*\*\*** | **0.28\*** | -0.11 | **-0.69\*\*\*** | 0.21 | 0.2 | 1178.47 | 1203.11 |
| Posterior cingulate | **0.22\*** | **-0.16\*\*** | -<br>**0.34\*\*** | **-0.28\*** | 0.26 | 0.09 | 0.08 | 1242.96 | 1267.61 |
| Precuneus | 0.13 | -0.1 | -0.21 | -0.04 | -0.07 | 0.02 | 0.02 | 1274.35 | 1298.99 |
| Putamen | -0.06 | **0.3\*\*\*** | **0.31\*\*** | -0.05 | **-0.54\*\*** | 0.15 | 0.14 | 1211.44 | 1236.08 |
| Superior frontal | 0.05 | -0.04 | 0.15 | **-0.35\*** | -0.09 | 0.03 | 0.02 | 1271.02 | 1295.66 |
| Superior parietal | 0 | 0.06 | -0.15 | 0.15 | 0.19 | 0.02 | 0.01 | 1278.06 | 1302.71 |
| Thalamus | 0.07 | 0.08 | 0.05 | -0.02 | **-0.63\*\*\*** | 0.05 | 0.04 | 1261.68 | 1286.32 |

\*\*\*\* $p < 0.001$ ; \*\* $p < 0.01$ ; \* $p < 0.05$ . **Abbreviations:** AIC, Akaike Information Criterion; BIC, Bayesian Information Criterion.

**Fig. 3.**
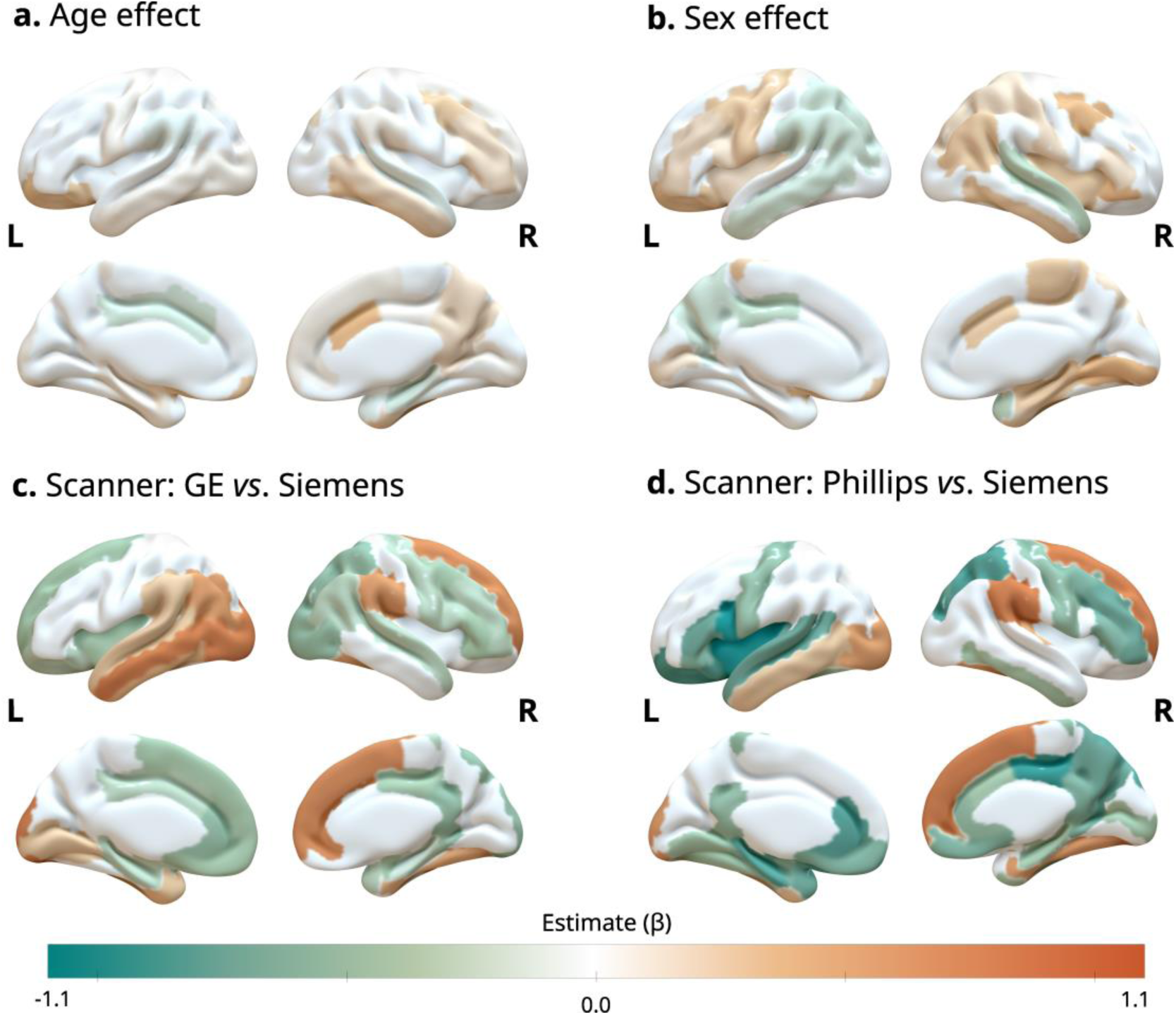
Cortical surface representation of standardized regression coefficients (β) for age, sex (males), and scanner-related predictors. A. Age effects; B. sex effects (males); C. GE *vs.* Siemens scanner effects; D. Philips *vs.* Siemens scanner effects. Only FDR-corrected significant associations (p < 0.05) are displayed.

Age-related associations were modest but spatially specific, affecting a limited set of cortical and subcortical regions. The strongest positive age effects were observed in subcortical nuclei, including the caudate, pallidum, and putamen bilaterally (β ≈ +0.30 to +0.39, *p < 0.001*), as well as in the hippocampus (β ≈ +0.12 to +0.13, *p < 0.05*). In contrast, age-related reductions in Z-score were primarily confined to posterior association cortices, including the posterior cingulate cortex (right β = −0.12, p < 0.05; left β = −0.16, *p < 0.01*) and angular gyrus bilaterally (β ≈ +0.22–0.23, *p < 0.001*, indicating relative preservation compared with reference regions). No significant age effects were observed in primary visual or parietal association cortices.

Sex-related effects were regionally restricted and of smaller magnitude. Significant associations were mainly observed in frontal and striatal regions, including the pallidum (right β = +0.34, *p < 0.01*; left β = +0.28, *p < 0.05*) and putamen bilaterally (β ≈ +0.31, *p < 0.05*). Outside frontal and basal ganglia regions, sex contributed minimally to regional Z-score variance.

The scanner manufacturer showed the most widespread and consistent regional effects. Frontal regions, particularly the superior frontal and medial orbitofrontal cortices, exhibited significantly lower Z-score values on GE (*e.g.*, right superior frontal: β = −0.46, *p < 0.001*) and Philips (*e.g.*, left medial orbitofrontal: β = −0.62, *p < 0.001*) scanners, relative to Siemens systems. Medial temporal structures were also strongly affected, with marked reductions in hippocampal Z-scores on both GE and Philips scanners bilaterally (β values ranging from −0.28 to −0.85; *p < 0.05* to *p < 0.001*). The occipital and parietal association cortices were less sensitive to biological variables but showed scanner-specific effects in selected regions.

### Scanner Technology Groups

Given the dominant and widespread effect of the scanner manufacturer, we investigated whether grouping scanners by technological generation would reduce this variability. Grouping by technology (stand-alone PET and PET/CT with or without time-of-flight) modestly reduced between-system variance. Still, meaningful differences persisted across groups, confirming that acquisition platform characteristics continue to influence Z-scores independently of the manufacturer label (Supplementary Figure 1-4).

### Cross-validation

In-sample model explanatory power was modest (R² = 0.01–0.23), consistent with the expectation that Z-score reflects considerable individual biological variability not captured by biological covariates. To assess the generalizability of each predictor’s predictive contribution, incremental variance (ΔR²) was computed using 10-fold cross-validation. Age and scanner manufacturer were the only variables showing meaningful out-of-sample contributions. Age produced a mean ΔR² of approximately 0.11 (range 0.02– 0.18), indicating a widespread and strong impact on Z-score across cortical and subcortical regions. Scanner manufacturers also contributed meaningfully, with a mean ΔR² of about 0.04 (range 0–0.10), consistent with systematic scanner-dependent variability. In contrast, sex contributed negligibly out-of-sample (mean ΔR² < 0.001). Full ΔR² distributions are shown in Supplementary Figure 5.

### Sensitivity analysis

To evaluate whether the inclusion of amyloid-positive individuals influenced the observed associations, all analyses were repeated after restricting the sample to amyloid-negative participants at baseline. A total of 278 participants were classified as amyloid-negative and included in the sensitivity analysis. Figure 4 demonstrates that normalization using the whole cerebellum and pons continued to produce widespread negative age-related associations across cortical regions, particularly in frontal and temporal areas. In contrast, cortical gray matter normalization again showed substantially fewer significant biological effects and reduced spatial heterogeneity.

**Fig. 4.**
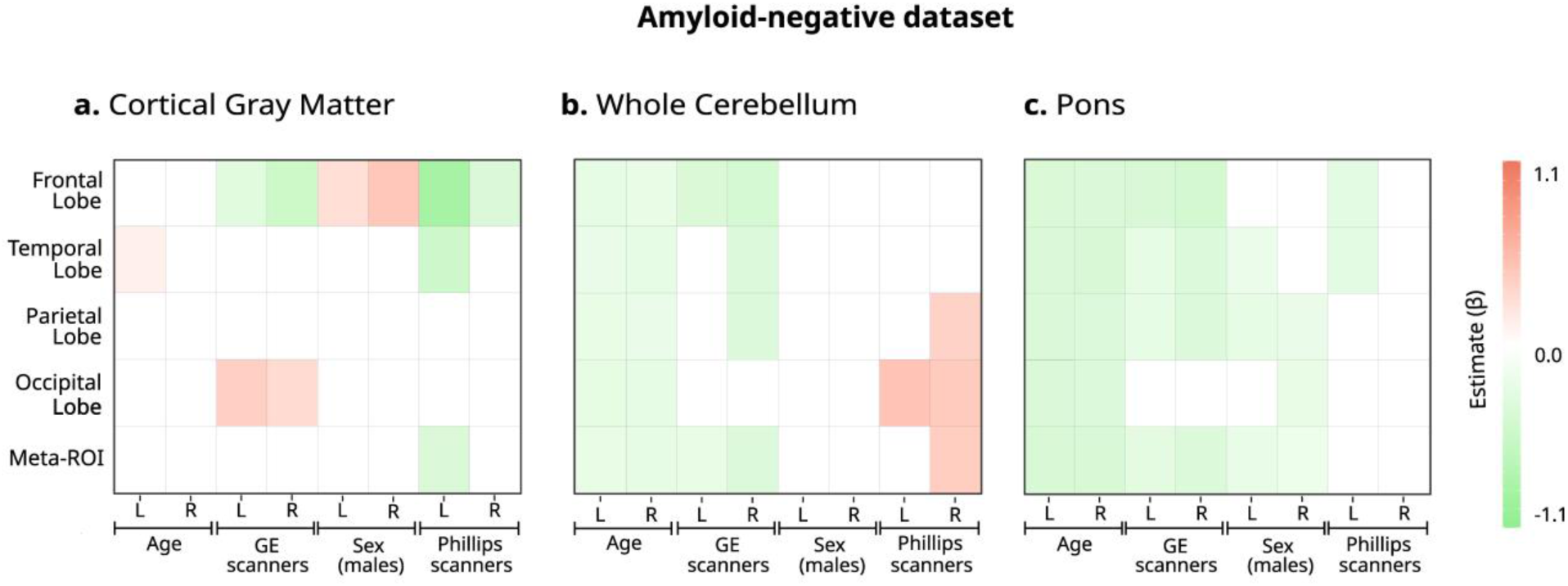
Heatmap of standardized regression coefficients (β) of biological and scanner-related predictors on lobar and AD meta-ROI [¹⁸F]FDG Z-scores, stratified by hemisphere, only for individuals that are amyloid-negative. A. Represents normalization by cortical gray matter; B. Represents normalization by whole cerebellum; C. Represents normalization by pons. Only coefficients with *p < 0.05* (FDR-corrected) are shown; nonsignificant associations are omitted to highlight the spatial distribution of significant effects.

Compared with the full sample (*n=449*), some associations became less spatially extensive and of lower statistical significance in the amyloid-negative subgroup. Quantitatively, the magnitude of scanner manufacturer effects decreased modestly: for example, the Philips-associated reduction in the left hippocampus decreased from β = −0.85 (full sample) to β = −0.65 (amyloid-negative), while the left medial orbitofrontal cortex decreased from β = −0.62 to β = −0.48. Sex-related effects were attenuated further and remained primarily restricted to frontal and subcortical regions. Scanner manufacturer effects persisted across all normalization strategies, confirming that technical artifacts are not driven solely by amyloid-positive individuals. Complete regional results of amyloid-negative estimates are provided in Supplementary Tables 5-7.

### Normative values

To provide a practical resource that accounts for these key confounders, we generated a covariate-stratified normative SUVr value for all brain regions. These were derived using the optimal normalization strategy identified, the cortical gray matter reference. Figure 5 presents the raw SUVr distributions that underlie our normative data, stratified by age and scanner manufacturer. While our primary analyses utilized Z-scores to facilitate cross-regional comparison, we present raw SUVr values here to provide clinically interpretable reference data. This visualization highlights the frontal lobe’s sensitivity to scanner effects. The complete set of normative data for all 83 brain regions and the meta-ROI, stratified by age, group, and scanner manufacturer, is available upon request.

**Fig. 5.**
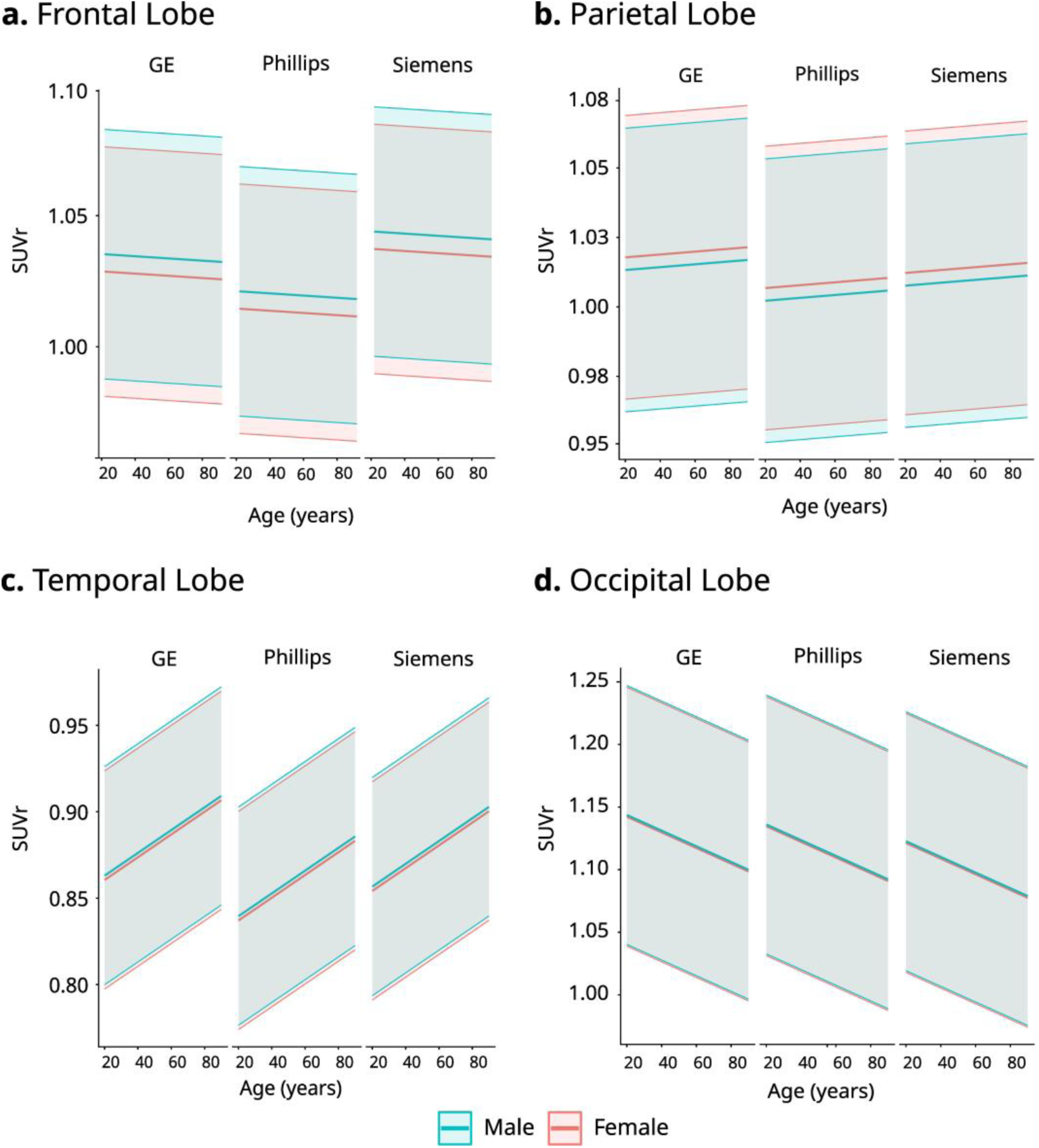
Normative [¹⁸F]FDG SUVr distributions across age, stratified by scanner manufacturer and sex. A. Frontal lobe; B. Parietal lobe; C. Temporal lobe; D. Occipital lobe. Lines represent mean SUVr for each age–manufacturer stratum, and shaded areas represent 95% confidence intervals.

## Discussion

Our key finding is that the scanner manufacturer significantly impacts regional Z-scores. This influence surpasses that of biological factors in both strength and area. Frontal lobe and medial temporal regions showed consistently lower Z-scores on GE and Philips scanners than on Siemens scanners, with effects reaching nearly one full standard deviation in some regions (*e.g.*, left hippocampus β = -0.85 for Philips). This region-specific bias could have direct clinical relevance, as analyzing scans from one manufacturer using normative databases or software calibrated for another manufacturer may lead to spurious classification of frontal metabolism. These systematic differences align with known vendor-specific variations in detector technology, attenuation correction, and reconstruction algorithms, which introduce intrinsic biases that were not eliminated by protocol standardization alone [26,27]. Attempts to reduce variability by grouping scanners into broader technological classes (*e.g.*, by PET/CT generation or TOF capability) yielded only modest improvements. This underscores that manufacturer identity remains a critical, irreducible source of variance in multicenter studies and highlights the urgent need for harmonization strategies that go beyond standardization of acquisition protocols. Advanced post hoc statistical harmonization methods, such as ComBat [28–30] or machine-learning-based techniques [31], have shown promise in mitigating these effects in neuroimaging data. These approaches should be considered a necessary step before pooling data from different vendors for normative modeling or machine-learning applications.

Age-related effects were robust, spatially structured, and biologically plausible. Positive age associations were observed not only in subcortical nuclei (*e.g.*, pallidum, putamen) but also in medial temporal lobe structures (hippocampus β ≈ +0.12, *p<0.05*), suggesting a pattern of relative metabolic preservation in these regions with normal aging. While these could reflect relative metabolic preservation, several observations suggest they may instead represent methodological artifacts. First, as shown in Figure 1, normalization to pons or whole cerebellum, regions known to decline with age, produces widespread negative age associations across all cortical lobes [19]. Second, even with cortical gray matter normalization, which minimizes this artifact, positive subcortical effects persist [32]. This likely occurs because the cortical gray matter reference region itself undergoes age-related decline, inflating target-to-reference ratios in regions with slower rates of decline [33]. Thus, the observed positive age effects should not be interpreted as true metabolic increases, but rather as a consequence of differential aging trajectories between reference and target regions. These complexities reinforce that age is a non-negotiable covariate in normative models, as confirmed by cross-validation (mean ΔR² ≈ 0.11 and ΔR² ≈ 0.04, respectively).

In contrast, the predictive contribution of sex was negligible out-of-sample (mean ΔR² < 0.001). This aligns with the observation that sex effects, although statistically significant in some frontal and striatal regions (*e.g.*, orbitofrontal cortex, pallidum), were small in magnitude (|β| < 0.37) and did not generalize to out-of-sample data. This confirms that while sex may be included for completeness in normative models, omitting it is unlikely to compromise their predictive accuracy in older CN populations, thereby simplifying model development.

A key methodological insight from our study is that the choice of reference region critically modulates sensitivity to biological and technical confounders. Normalization to the pons or the whole cerebellum introduced strong, widespread age-associated effects. In contrast, normalization to cortical gray matter consistently minimized the influence of age, sex, and scanner manufacturer, yielding the most stable and internally consistent Z-scores. This finding is crucial for practice and recommends cortical gray matter as the preferred reference for studies aiming to isolate disease-specific metabolic changes from normal variation and scanner artifacts. Furthermore, the AD meta-ROI exhibited robustness, with minimal variance explained by any confounder. This supports the use of such composite measures as stable biomarkers, particularly in multisite research where technical variability is a concern [23].

Our sensitivity analysis with amyloid-negative participants showed that the spatial patterns of age-, sex-, and scanner-related effects were consistent with those in the full cohort. The magnitude of the scanner manufacturer effect was modestly reduced; for example, the Philips-associated left hippocampus decreased from β = −0.85 to β = −0.65. This attenuation might be due to three mechanisms. First, amyloid-positive CN individuals may show early hypometabolic changes. Their exclusion, potentially unevenly distributed across scanners and more frequent on Philips, reduces the scanner effect. Second, the smaller amyloid-negative sample (278 *vs*. 449 individuals) decreases power and attenuates extreme coefficient estimates (βs). Third, amyloid-positive cases may have more age-related atrophy, which could amplify manufacturer differences. Despite these factors, scanner effects remained significant and clinically relevant.

This study has several limitations. First, the majority of scans were acquired on older-generation analog PET systems (e.g., GE Discovery ST, Siemens Biograph). Given the distinct noise and contrast recovery characteristics of modern digital detectors and time-of-flight systems, our vendor-specific effect sizes should not be directly extrapolated to contemporary scanners without replication. Second, the distribution of scanners across manufacturers was unequal, with Siemens systems overrepresented relative to GE and Philips. This imbalance may reduce the precision of manufacturer-specific estimates and limit the generalizability of our findings. Third, SUVr remains a semi-quantitative measure that does not capture the full physiological specificity available with fully quantitative techniques (*e.g.*, absolute cerebral metabolic rate of glucose, CMRglc). Fourth, the small sample sizes at the extremes of the age spectrum (particularly participants under 65 years, n=21) necessitated broad age categories, which may obscure more detailed nonlinear age trajectories. Fifth, another potential source of variability is the partial volume effect resulting from cortical atrophy. Age-related gray matter loss can artificially reduce PET signal in cortical regions, potentially confounding metabolic estimates. Because the present study relied on SUVr measures without explicit partial volume correction, part of the observed age effects may reflect structural rather than purely metabolic differences. Finally, all scanner-related conclusions are based on a cross-sectional comparison between groups of participants; ideally, a head-to-head acquisition of the same subjects on different scanners would be required to fully isolate vendor-specific effects, though this is rarely feasible in large cohorts.

## Conclusion

This study demonstrates that both technical and biological factors influence normative quantification of [¹⁸F]FDG-PET in CN older adults, with scanner manufacturer and age emerging as the most relevant predictors of regional Z-score variability, while sex showed minimal out-of-sample predictive value. Scanner-related differences were particularly pronounced in frontal and medial temporal regions, where GE and Philips systems consistently produced lower Z-scores than Siemens scanners, with effects approaching one standard deviation. Our findings also show that the choice of reference region substantially modulates these confounding effects, with cortical gray matter normalization providing the most stable results and minimizing age and scanner-related variability compared with pons or whole cerebellum references. In addition, composite measures such as the AD meta-ROI demonstrated high robustness to both technical and biological variability, supporting their use as reliable biomarkers. These results highlight the importance of considering scanner features, age, and normalization when interpreting [¹⁸F]FDG-PET data. They offer stratified normative SUVr and Z-score values to enhance reliability and comparability of metabolic PET analyses in aging groups.

## Acknowledgments

Data collection and sharing for this project was funded by the Alzheimer’s Disease Neuroimaging Initiative (ADNI) (National Institutes of Health Grant U01 AG024904) and by the Department of Defense ADNI (award number W81XWH-12-2-0012). ADNI is funded by the National Institute on Aging, the National Institute of Biomedical Imaging and Bioengineering, and through generous contributions from the following: AbbVie, Alzheimer’s Association; Alzheimer’s Drug Discovery Foundation; Araclon Biotech; BioClinica, Inc.; Biogen; Bristol-Myers Squibb Company; CereSpir, Inc.; Eisai Inc.; Elan Pharmaceuticals, Inc.; Eli Lilly and Company; EuroImmun; F. Hoffmann-La Roche Ltd and its affiliated company Genentech, Inc.; Fujirebio; GE Healthcare; IXICO Ltd.; Janssen Alzheimer Immunotherapy Research & Development, LLC.; Johnson & Johnson Pharmaceutical Research & Development LLC.; Lumosity; Lundbeck; Merck & Co., Inc.; Meso Scale Diagnostics, LLC.; NeuroRx Research; Neurotrack Technologies; Novartis Pharmaceuticals Corporation; Pfizer Inc.; Piramal Imaging; Servier; Takeda Pharmaceutical Company; and Transition Therapeutics.

The Canadian Institutes of Health Research is providing funds to support ADNI clinical sites in Canada. Private sector contributions are facilitated by the Foundation for the National Institutes of Health (www.fnih.org). The grantee organization is the Northern California Institute for Research and Education, and the study is coordinated by the Alzheimer’s Therapeutic Research Institute at the University of Southern California. ADNI data are disseminated by the Laboratory for Neuro Imaging at the University of Southern California.

## Statements & Declarations

### Author Contributions

Giordana Salvi de Souza: Conceptualization, Formal analysis, Visualization, Writing – Original Draft, Writing – Review & Editing; Guilherme Povala: Conceptualization, Methodology, Formal analysis, Validation, Writing – Review & Editing; Guilherme Garcia Schu Peixoto: Conceptualization, Methodology, Formal analysis, Validation, Writing – Review & Editing; Artur Martins Coutinho: Methodology, Writing – Review & Editing; Andrei Bieger: Data curation, Writing – Review & Editing; Mateus Rozalem-Aranha: Methodology, Writing – Review & Editing; Marco Antônio de Bastiani: Formal analysis, Writing – Review & Editing; Eduardo Rigon Zimmer: Writing – Review & Editing; Wyllians Vendramini Borelli: Conceptualization, Writing – Review & Editing; Laura Willers de Souza: Project administration, *Supervision,* Writing – Review & Editing.

### Data Availability

Data used in preparation of this article were obtained from the Alzheimer’s Disease Neuroimaging Initiative (ADNI) database (adni.loni.usc.edu).

### Ethics Approval

This study was conducted in accordance with the ethical standards of the institutional and/or national research committee and with the 1964 Declaration of Helsinki and its later amendments, or comparable ethical standards. Data used in this study were obtained from the Alzheimer’s Disease Neuroimaging Initiative (ADNI) database. The ADNI study was approved by the institutional review boards of all participating sites.

### Consent to Participate

Informed consent was obtained from all individual participants included in the study.

### Consent to Publish

Not applicable. This manuscript does not contain any individual person’s data in any form (including any individual details, images, or videos).

