## Supplementary_Material for "Sources of Variability in Normative Cerebral [^18^F]FDG-PET imaging"

**Supplementary Methods: Cross-validation and Incremental R² Analysis**

Regional FDG-PET standardized uptake value ratios (SUVr) were available for 86 brain regions per participant. Candidate covariates included age, sex, and scanner manufacturer. Analyses were conducted on the full cohort after standard quality control procedures.

#### ***Statistical modeling***

For each brain region, we fitted the following linear model using ordinary least squares:

SUVr_region_ ∼ sex + age + scanner_manufacturer

Generalization performance and variable importance were evaluated using 10-fold cross-validation. Folds were defined at the participant level (RID) to ensure that no individual contributed to both training and test sets within a fold. Numeric predictors were centered and scaled using statistics from the training fold, which were then applied to the held-out fold to avoid data leakage. The same fold structure was reused across regions to ensure comparability of out-of-fold (OOF) predictions.

**Incremental predictive contribution**

To quantify each variable’s unique predictive contribution, we computed the cross-validated incremental R² for each predictor *p* and region:

ΔR_p_^2^ = R_full_^2^ − R_reduced(−p)_^2^,

where R_full_^2^ is the OOF R^2^ from the full model and R_reduced(−p)_^2^ from the same model omitting *p*. We summarize ΔR² per region and predictor by its fold-wise mean (and SD). Because ΔR² is computed from out-of-sample predictions, it provides a measure of predictive importance rather than statistical significance. This approach reduces inflation due to overfitting and better reflects each variable’s generalizable contribution to PET signal variability. As a practical criterion for covariate retention, we defined ΔR² ≥ 0.01 (≥1% OOF variance explained) as a meaningful contribution.

#### **Results and Interpretation**

Across the 86 regions, age showed the largest and most consistent contribution (mean ΔR² ≈ 0.11; range 0.02–0.18), indicating a strong effect of aging on brain glucose metabolism. Scanner manufacturer showed a moderate contribution (mean ΔR² ≈ 0.04; range 0–0.10), consistent with systematic differences between acquisition systems.

By contrast, sex contributed negligibly to predictive accuracy (mean ΔR² ≈ 0.0008).

#### **Summary**

These results indicate that age and scanner manufacturer meaningfully improve out-of-sample prediction of FDG-PET SUVr across brain regions and should therefore be retained as covariates in normative modeling. In contrast, sex provides negligible predictive value and can be omitted without loss of performance.

### **Supplementary Figures**

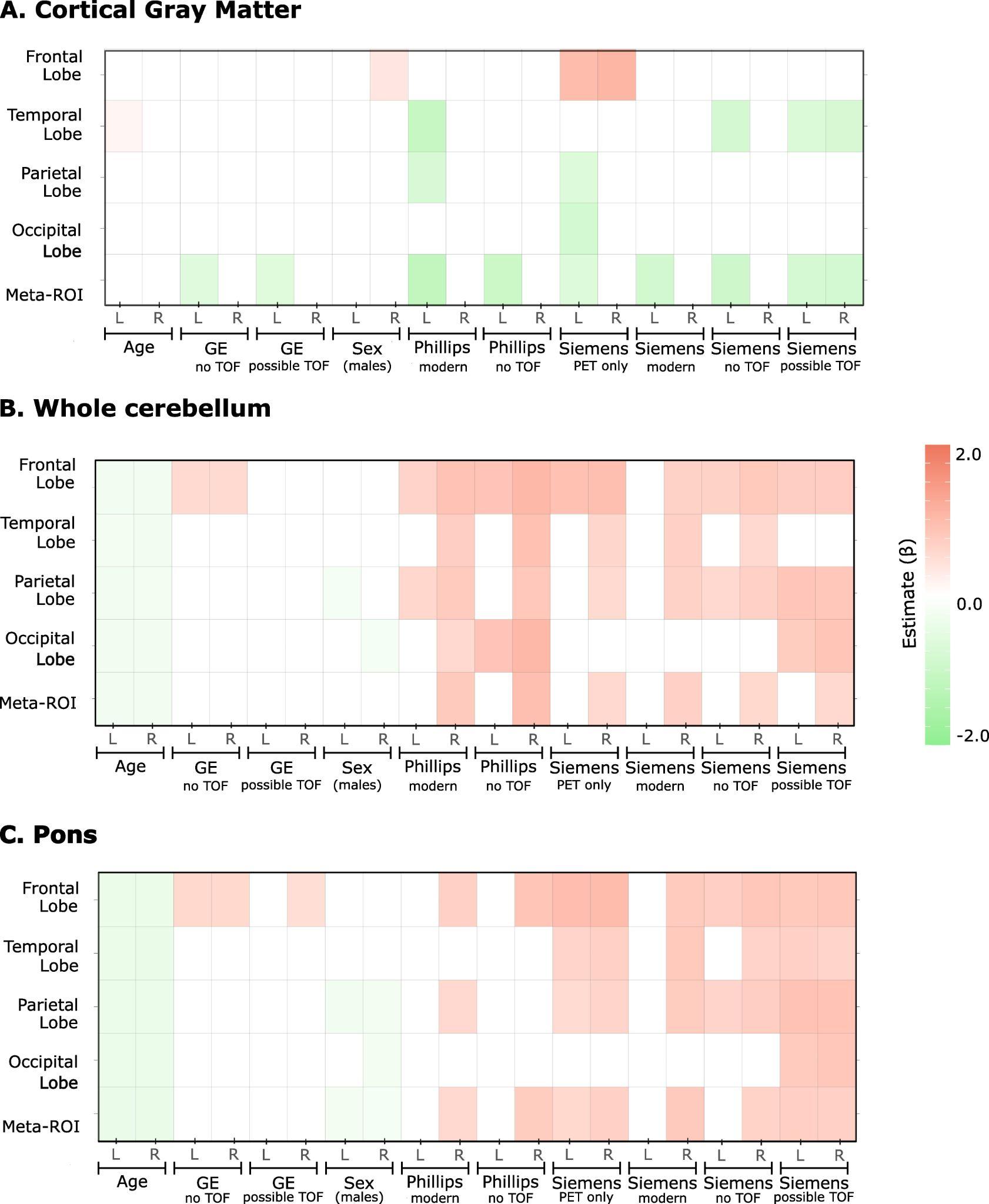

**Supplementary Figure 1:** Predictors of [^18^F]FDG Z-score for lobes and meta-ROI using three different regions of reference. Only significant estimates (*p<0.05*) are presented in the heatmap.

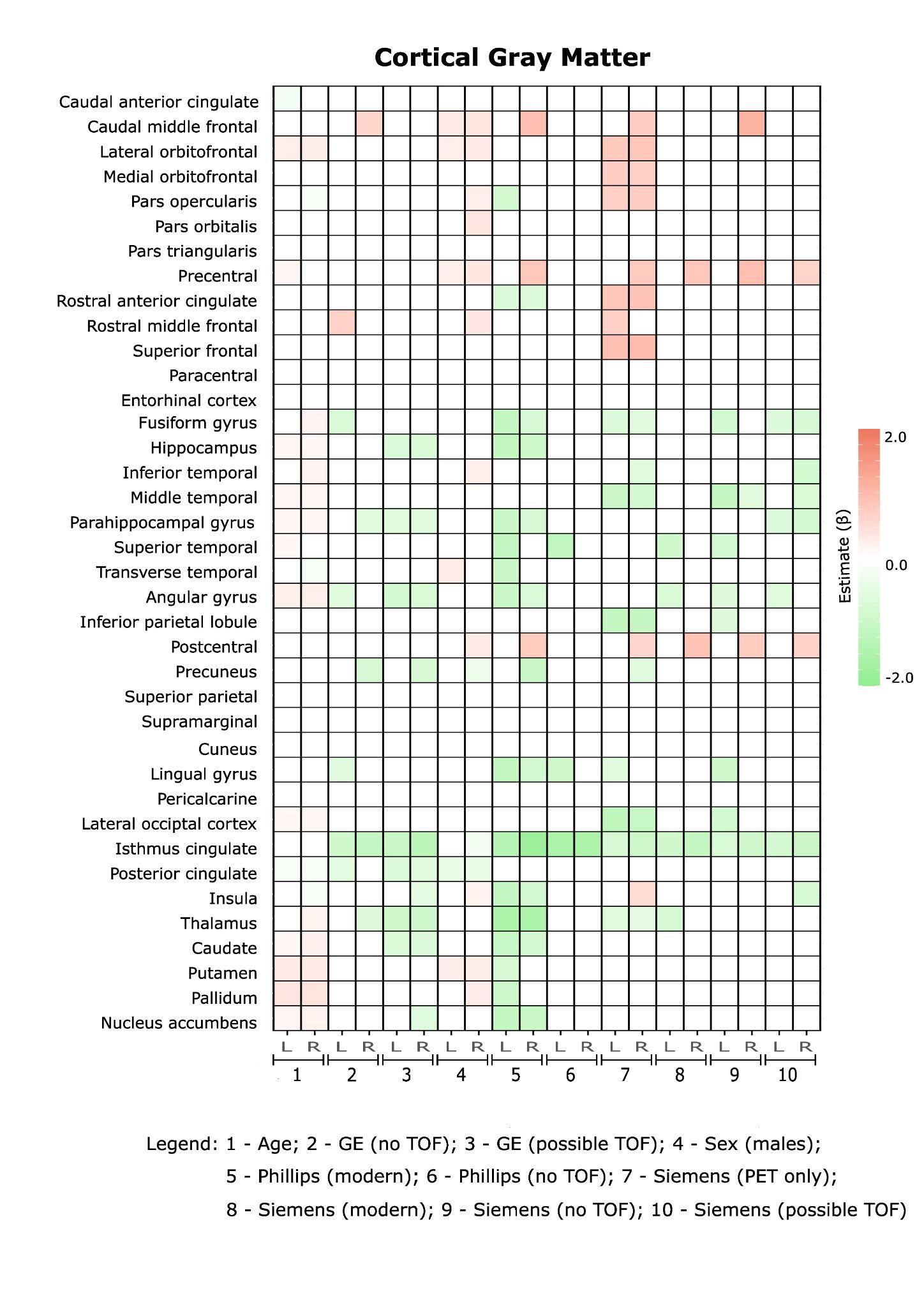

**Supplementary Figure 2:** Predictors of [^18^F]FDG Z-score for all brain regions using cortical gray matter as region of reference. Only significant estimates (*p<0.05*) are presented in the heatmap.

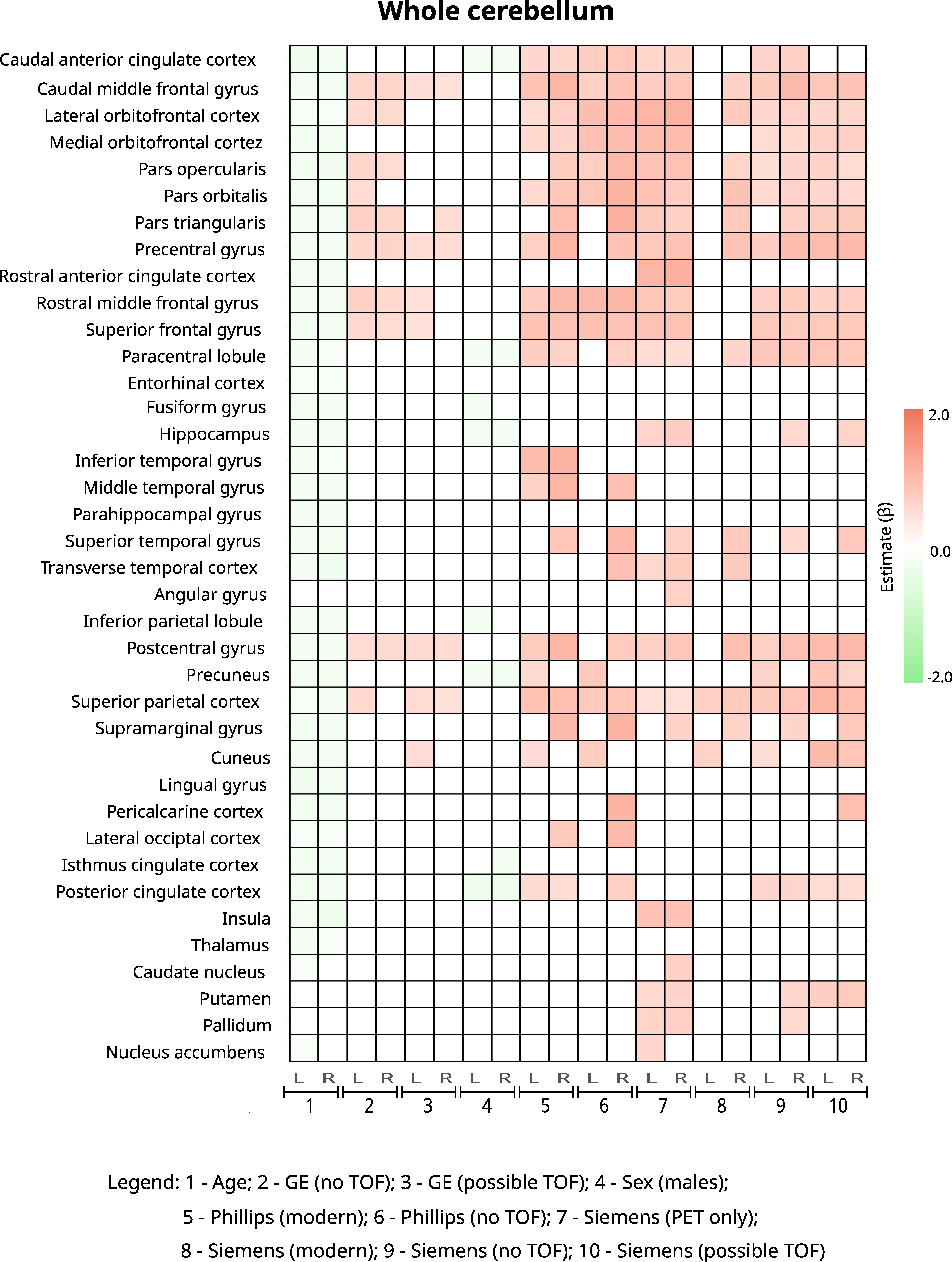

**Supplementary Figure 3:** Predictors of [^18^F]FDG Z-score for all brain regions using whole cerebellum as the region of reference. Only significant estimates (*p<0.05*) are presented in the heatmap.

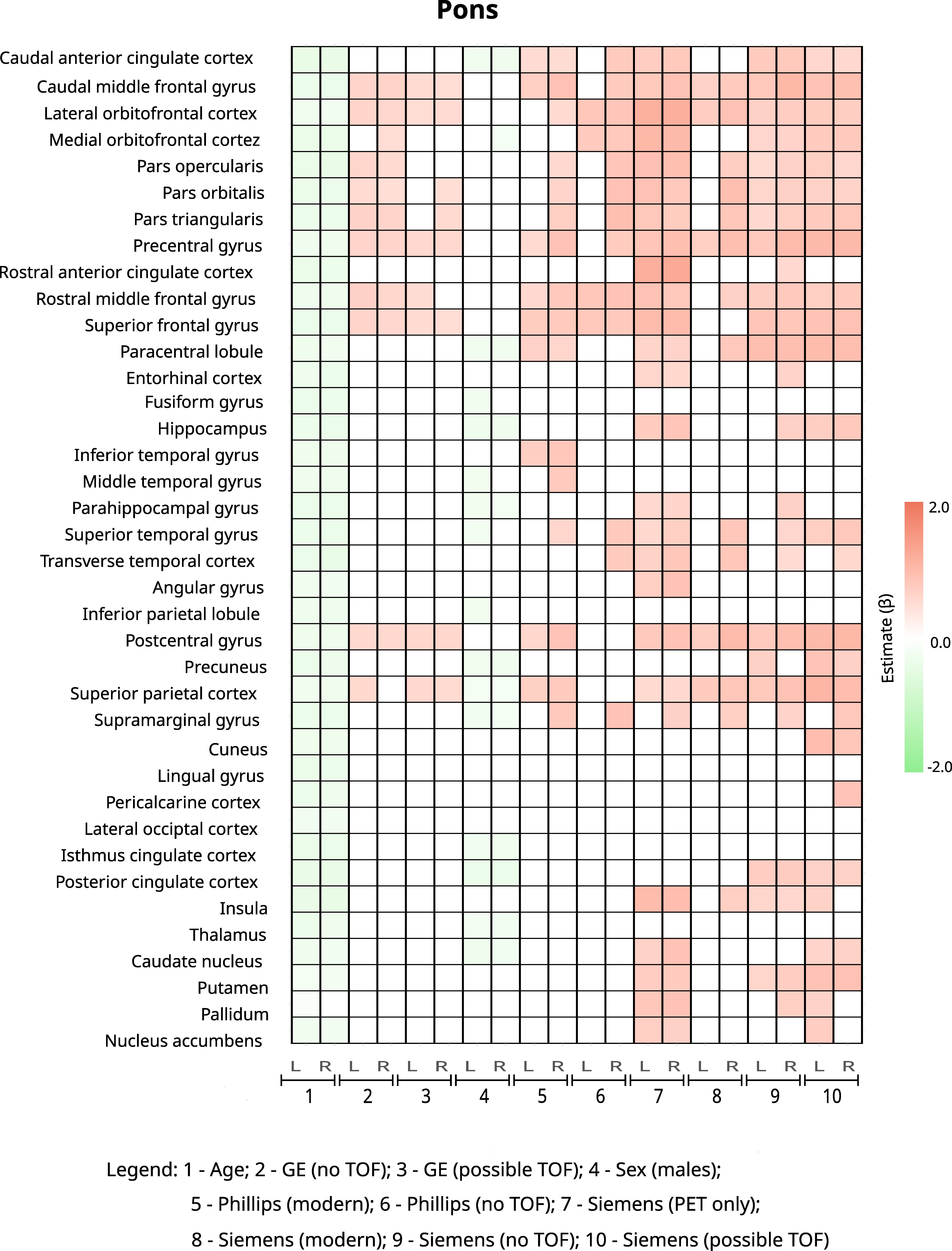

**Supplementary Figure 4:** Predictors of [^18^F]FDG Z-score for all brain regions using pons as region of reference. Only significant estimates (*p<0.05*) are presented in the heatmap.

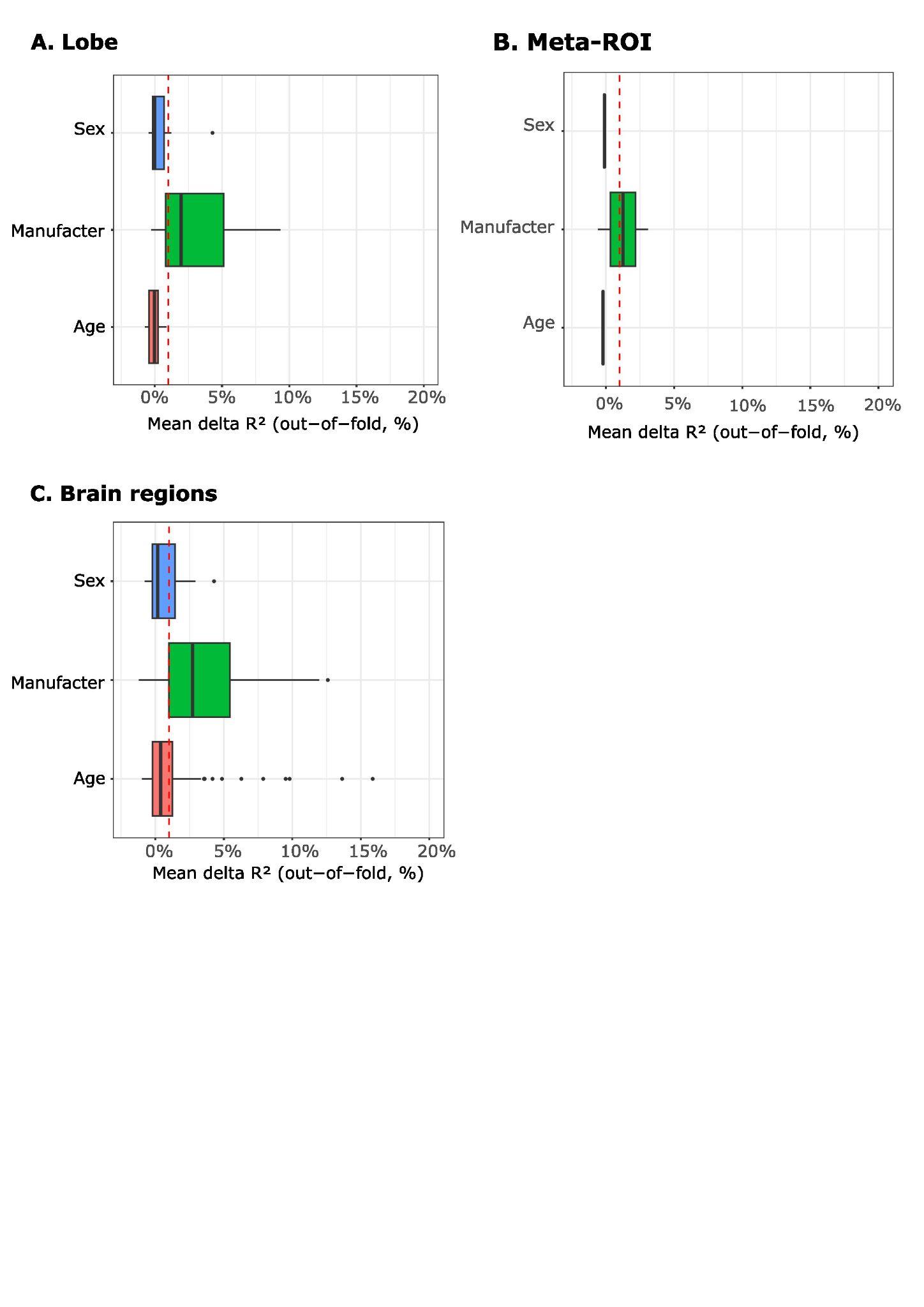
**Supplementary Figure 5:** Cross-validation results for cortical gray matter as reference region.
